# Repeated Caffeine Intake Suppresses Cerebral Grey Matter Responses to Chronic Sleep Restriction in an A_1_ Adenosine Receptor-Dependent Manner

**DOI:** 10.1101/2023.07.23.550201

**Authors:** Yu-Shiuan Lin, Denise Lange, Diego Baur, Anna Foerges, Congying Chu, Changhong Li, Eva-Maria Elmenhorst, Bernd Neumaier, Andreas Bauer, Daniel Aeschbach, Hans-Peter Landolt, David Elmenhorst

**Affiliations:** Centre for Chronobiology, Psychiatric Hospital of the University of Basel, Basel, Switzerland; Research Platform Molecular and Cognitive Neurosciences, University of Basel, Basel, Switzerland; Athinoula A. Martinos Center for Biomedical Imaging, Mass General Hospital, Harvard Medical School, Boston, U. S. A; Department of Sleep and Human Factors, Institute of Aerospace Medicine, German Aerospace Center, Cologne, Germany; Institute of Pharmacology and Toxicology, University of Zurich, Zurich, Switzerland; Sleep & Health Zurich, University Center of Competence, University of Zurich, Zurich, Switzerland; Institute of Neuroscience and Medicine, INM-2, Forschungszentrum Jülich, Jülich, Germany; Department of Neurophysiology, Institute of Zoology (Bio-II), RWTH Aachen University, Aachen, Germany; Institute for Occupational, Social, and Environmental Medicine, RWTH Aachen University, Aachen, Germany; Institute of Neuroscience and Medicine, INM-5, Forschungszentrum Jülich, Jülich, Germany; Institute of Experimental Epileptology and Cognition Research, University of Bonn Medical Center, Bonn, Germany; Multimodal Neuroimaging Group, Department of Nuclear Medicine, University Hospital Cologne, Cologne, Germany

## Abstract

Evidence has shown that both sleep loss and daily caffeine intake can induce changes in grey matter (GM). Caffeine is frequently used to combat sleepiness and impaired performance caused by insufficient sleep. It is unclear 1) whether *daily* use of caffeine could prevent or exacerbate the GM alterations induced by *chronic* sleep restriction, and 2) whether the potential impact on GM plasticity depends on individual differences in the availability of adenosine receptors, which are involved in mediating effects of caffeine on sleep and waking function. In this double-blind, randomized, controlled study, 36 healthy adults (aged 28.9 ± 5.2 y/o; 15 females; habitual daily caffeine intake < 450 mg; 29 homozygous C/C allele carriers of the A2A adenosine receptor (A_2A_R) gene variant rs5751876 of *ADORA2A*) underwent a 9-day laboratory visit consisting of one adaption day, 2 baseline days (BL), 5-day sleep restriction (CSR, 5 h time-in-bed), and a recovery day (REC) after an 8-h sleep opportunity. Nineteen participants received 300 mg caffeine in coffee through the 5 days of CSR (CAFF group), while 17 matched participants received decaffeinated coffee (DECAF group). We measured the GM morphology on the 2^nd^ BL Day, 5^th^ CSR Day, and REC Day. Moreover, we used [^18^F]-CPFPX PET to quantify the baseline availability of A_1_ adenosine receptors (A_1_R) and their relation to GM plasticity. The voxel-wise multimodal whole-brain analysis on T1-weighted images controlled for variances of cerebral blood flow indicated a significant interaction between caffeine and CSR in four brain regions: 1) right temporal-occipital region, 2) right thalamus, 3) left dorsolateral, and 4) dorsomedial prefrontal region. The post-hoc analyses indicated increased GM intensity in the DECAF group in all four regions but decreased GM in the thalamus as well as dorsolateral and dorsomedial prefrontal regions in the CAFF group after sleep restriction. Furthermore, lower baseline subcortical A_1_R availability predicted larger reduction in the CAFF group after CSR of all brain regions except for the caffeine-associated thalamic reduction. In conclusion, our data suggest an adaptive upregulation in GM after 5-day CSR, while concomitant use of caffeine instead leads to a GM reduction. The lack of consistent association with individual A_1_R availability may suggest that CSR and caffeine affect GM plasticity predominantly by a different mechanism. Future studies on the role of adenosine A_2A_ receptors (ADORA2A) in CSR-induced GM plasticity are warranted.

## Introduction

Caffeine is the most widely used psychoactive substance (1). Given its efficacy in improving alertness (2) and alleviating cognitive impairments caused by sleep deprivation (3) or sleep restriction (4), it is often consumed to combat drowsiness (5). On the cerebral level, both acute sleep loss and daily caffeine intake can lead to a reduction in human grey matter (GM) volumes (6–10) as measured by Magnetic Resonance Imaging (MRI). It is unclear whether use of caffeine alleviates or exacerbates the sleep loss associated changes in GM after insufficient sleep.

Impaired brain structures and functionality have been frequently found after sleep deprivation (11) and sleep restrictions (12–14) in rodents. In humans, one night of sleep restriction (i.e. measuring after a night of 3-h time in bed) leads to cortical changes specifically in a young age group, including decreased GM in the thalamus, precuneus, and postcentral gyrus but increased GM in the insula (6). One month of sleep restriction, on the other hand, resulted in a reduction in the white matter diffusivity (15). On the other hand, other studies reported several, albeit inconsistent, non-linear changes over time in multiple cortical regions from 24h to 72h of total sleep deprivation (7, 8). Effects of *chronic* sleep restriction on GM have not yet been investigated. Furthermore, chronic caffeine administration was found to inhibit hippocampal neurogenesis and cell proliferation in adult rodents (16–18), while *daily* caffeine intake was found to be associated with reduced GM volume in multiple cortical regions including hippocampus in humans (9, 10). We hypothesize that GM can be reduced by CSR, while taking caffeine during *chronic* sleep restriction can exacerbate the GM reduction to sleep restriction.

Effects of both caffeine and sleep loss are mediated at least in part by the adenosine system (19, 20). Partly serving as a byproduct of neuronal activity, the release of extracellular endogenous adenosine is increased throughout wakefulness (21), and the binding of adenosine to the A_1_ adenosine receptor (A_1_R) leads to neural inhibition. Extended wakefulness, including partial or complete sleep deprivation can lead to increased levels of extracellular adenosine (22) and upregulated A_1_R binding (23). On the other hand, caffeine enhances synaptic excitation by non-selective antagonism of adenosine at the adenosine receptors (24), thereby increasing alertness (19), reducing reaction time (2), and counteracting sleep loss-induced cognitive decline (3, 4). Furthermore, individual variabilities in the magnitude of cerebral and behavioral responses to caffeine and sleep loss is partly determined by the A_1_R availability (25–27). Hence, we expect that participants’ baseline A_1_R binding potentials may predict the magnitudes of GM plasticity induced by CSR and caffeine. In addition, since the sensitivity to caffeine effects on sleep (28) and the distribution of A_1_R (29) are known associate with the variant rs5751876 of the A_2A_ adenosine receptor encoding gene, *ADORA2A*, we genotype this gene variant in the study participants to address our research questions in groups with matched distribution in age, gender, body mass index, chrono type, and level of habitual caffeine intake (4).

As the vasoconstrictive effect of caffeine can affect morphometric measurements through altering perfusion (30), we quantify cerebral blood flow (CBF) as a covariate for the morphological analysis. As an exploratory step, we also examined whether CBF was altered after chronic sleep restriction, as earlier studies have found rather region-dependent mixed effects after different durations of sleep loss. A study reported a reduced absolute CBF (absCBF) in the attention network after *acute* sleep restriction (4h in bed) in subjects with higher drowsiness but elevated absCBF in basal forebrain in those who remained alert (31). Other studies employing one-night sleep deprivation have found both reduced relative CBF (rCBF) in parahippocampus, fusiform, and prefrontal cortices (32) as well as increased rCBF in occipital cortices and insula (33). Similarly, one month of sleep restriction (5.5h in bed) led to a reduced diffusivity in the orbitofrontal gyri, superior occipital gyri, insula, and fusiform but increased supplementary motor area and cingulate gyrus (15). How *chronic* sleep restriction affects perfusion, specifically CBF remains unknown and is explored additionally in this study.

## Methods

The data collection of this study took place at the: envihab research facility of the German Aerospace Center (Cologne, Germany), and the study protocol was approved by the Ethics Committee of the regional Medical Board (Ärztekammer Nordrhein) and the German Federal Office for Radiation Protection. All the participants have given informed consent in written form.

### Study procedure

In a double-blind randomized study, 36 healthy adults (aged 28.9 ± 5.2 y/o; 15 females; habitual daily caffeine intake < 450 mg; variant of ADORA2A rs5751876 allele: 29 C/C homozygous, 6 C/T heterozygous, 1 T/T homozygous; for detailed inclusion and exclusion criteria see (27)) underwent 9 consecutive laboratory days in the following order: 1-day adaptation (8-h time in bed), 2-day baseline (BL, 8-h time in bed/day), 5-day CSR days (5-h time in bed/day), 1-day recovery with 8h time in bed (REC; **Fig 1**). All participants followed a one-week sleep satiation protocol (9h in bed between 22:00 and 07:00 or between 23:00 and 08:00) prior to the start of the study. The compliance was monitored by daily self-reports and actimetry. Starting from the adaptation day, participants were in constant lighting (∼100 lx) control during wakefulness. Across the 5-day CSR, 19 of the 36 participants received daily coffee containing 200 mg and 100 mg of caffeine in the morning and at noon, respectively (CAFF group), while the other 17 participants (DECAF group) received decaffeinated coffee (4). [^18^F]-CPFPX PET-MRI acquisitions (Biograph mMR, Siemens) took place in the afternoon of the second BL Day, the 5^th^ CSR Day (average 7.0 ± 0.8h after the last caffeine intake), and the REC Day (average 29.8 ± 4.3h after the last caffeine intake). Salivary samples were collected repeatedly every evening, except for the adaptation day, as well as in the morning of the baseline days and before the PET scans (i.e., 1^st^ and 2^nd^ BL Day, 5^th^ day of CSR and REC Day) to measure the caffeine concentrations.

**Figure 1.**
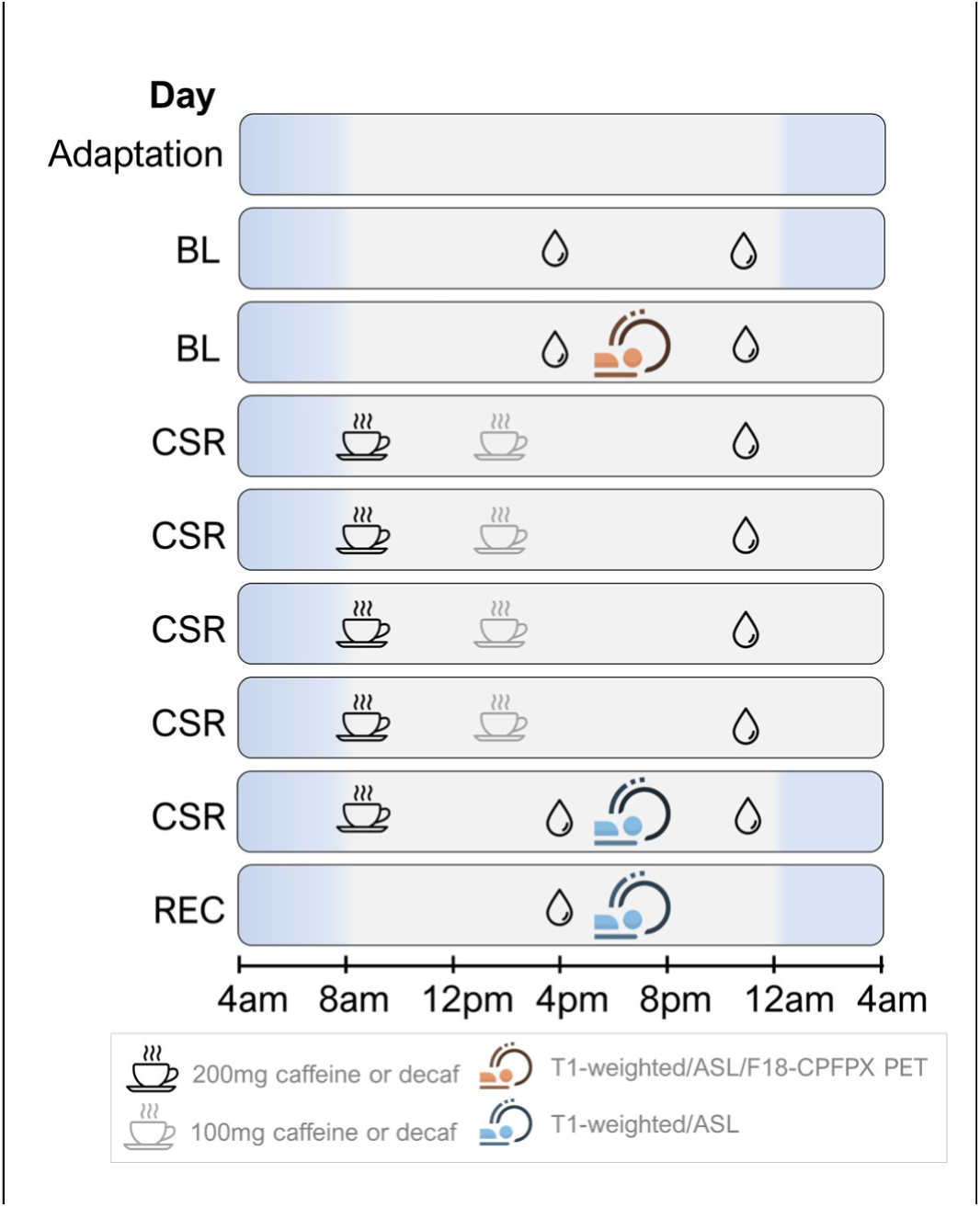
An overview of study protocol.

### Data acquisition and processing

Overall, we first identified the cortical changes in response to the main effect of CSR and the interaction of Caffeine x CSR. We used whole-brain voxel-wise multimodal linear mixed models, which allowed regressing out the variance of CBF voxel-to-voxel with Arterial Spin Labeling (ASL) images. Next, we conducted an independent whole-brain analysis on ASL images to determine the main effect of CSR and the interaction of Caffeine x CSR on CBF. Lastly, we extracted the mean intensity of GM and mean CBF of the identified regions, as well as the binding potential (BP_ND_) of A_1_R in cortical and subcortical regions from all time points (i.e., baseline, CSR Day, and REC Day), respectively. Using the extracted values, we then conducted post-hoc analyses as well as examined the association between the GM and/or CBF responses and the A_1_R BP_ND_ with linear regressions.

#### T1-weighted images and GM intensity

Grey matter (GM) morphology was assessed by T1-weighted images using 3-dimensional magnetization-prepared rapid acquisition with gradient echo (3D MPRAGE; isotropic voxel 1mm, TR= 2250ms, TE= 3.03ms). For the preprocessing of T1-weighted images, we used the “Segment Longitudinal Data” pipeline provided by CAT12 toolbox on SPM12 (University College London, London, UK), which enabled a co-registration with the mean of the three volumes collected from the three timepoints (i.e. BL, CSR, and REC) of each participant and was suitable for repeated measurements. Affine registration was conducted using the tissue probability map in SPM12, followed by the segmentation of brain tissues into grey matter, white matter, cerebrospinal fluid, as well as total intracranial volume. The procedure continues with the modulated normalization using an MNI (Montreal Neurological Institute, McGill University) –defined standard brain. Lastly, we smoothed the preprocessed data with Gaussian kernel of FWHM= 8 x 8 x 8 mm^3^.

Next, we identified the GM responses to CSR and caffeination x CSR using a voxel-wise multimodal analysis in order to statistically control for the potential bias caused by caffeine-induced changes in brain perfusion (30, 34, 35). We used VoxelStats toolbox (36), with which we conducted linear mixed models on two types of co-registered 3D volumes, i.e., T1-weighted and ASL, through each corresponding voxel. In addition to CBF, we used age, sex, and total intracranial volume as regressors of no interests in the model. The statistical significance of the coefficients was acquired with permutation tests (5000 times) and defined by a threshold at cluster-level p_FDR_ < .001)

For the regions identified to be responsive to the interaction of Caffeine x CSR vs BL, we further conducted a post-hoc analysis to determine the exact GM changes in each group in response to CSR using region-of-interest-based approach on R (R core team, Vienna, Austria). We first created masks of the responsive regions identified in the whole-brain analyses. Accordingly, we extracted mean responses of these regions from all time points (BL, CSR, and REC) using FSL *fslmeants*, followed by a post-hoc analysis to determine the directions of GM responses in each group.

#### Cerebral Blood Flow

Cerebral blood flow (CBF) was measured by the sequence ASL (5×5×5mm^3^, TR =3500ms, TE= 11ms). For the preprocessing of ASL images, we used the standard pipeline from FMRIB Software Library (FSL 5.0; Oxford Center for Functional MRI of the Brain, United Kingdom). We used the first volume for the reference of motion correction as well as for the M0 calibration. We calculated the relative CBF maps with the acquired 40 tag-control pairs, followed by the quantification of absolute CBF using white matter as the reference tissue. Lastly, the absolute CBF maps were co-registered onto the T1-weighted images and MNI space.

Next, we conducted the whole-brain analysis on the absolute CBF maps with a GM mask using 2-way mixed effect ANOVA. Similar to the analysis on T1-weighted images, we also used nonparametric threshold-free cluster enhancement (number of permutations = 5000, cluster-level threshold p < .01) with the “*randomise*” function. Same as the GM analysis, a post-hoc analysis was carried out on the significant changes in response to Caffeination x CSR using an identical procedure in extracting the regional mean response and post-hoc analyses.

#### Cerebral A_1_R availability

We acquired the [^18^F]-CPFPX PET data (framing scheme: 4 × 60 s, 3 × 120 s, 18 × 300 s, 2.09 × 2.09 × 2.03 mm³) simultaneously with the magnetic resonance imaging (MRI) scan. The scan in total lasted 100 minutes. We adopted an intravenous bolus injection simultaneously with the start of the scan, followed by a constant infusion (15.9 ml/ 34.1 ml, Kbol = 55 min.) The eventual dose injected was 175.9 ± 21.8 MBq and the molar activity was 102.25 ± 72.08 GBq/µmol at the time of injection.

For the preprocessing of PET, we used the standard pipeline of PMOD Neuro Tool (v4.006; PMOD Technologies). We first conducted motion correction with the average of the PET data collected in the first 9 min. The matching PET image was co-registered with the corresponding preprocessed T1-weighted image and segmented into grey matter, white matter, and cerebrospinal fluid, followed by the spatial normalization based on the MNI space. Seventy volumes of interests (VOIs) were defined by the automated anatomical labelling template of MNI space. For the kinetic modeling we used PMOD Kinetics Tool (version 4.006; PMOD Technologies). We used cerebellum as a reference region to calculate the time-activity curves (TACs) of each volume of interest. The availability of A_1_R was indicated by the binding potential BP_ND_ of [^18^F]-CPFPX acquired with the Logan’s reference tissue model (t* = 30 min (37)).

We reported the details of procedure and parameters with regard to PET data acquisition, preprocessing, and kinetic modeling in the method section of (27).

#### Predicting cerebral responses by cortical and subcortical A_1_R BP_ND_

In a last step, we used A_1_R BP_ND_ to predict the identified cerebral responses (i.e., changes in GM and CBF). We first calculated the percent changes in CSR relative to BL using (meanCSR – meanBL)/meanBL * 100%. We calculated the A_1_R BP_ND_ data by cortical and subcortical regions. For cortical A_1_R BP_ND_, we averaged the values acquired in frontal, occipital, parietal, temporal, and cingulate cortices, as well as amygdala, hippocampus, and insula. For subcortical A_1_R BP_ND_, we averaged the values acquired in pallidum, olfactory tubercle, caudate nucleus, putamen, and thalamus by each hemisphere. The regions were segmented based on the automated anatomical labeling atlas (AAL (38)). We then conducted linear regression model to examine the coefficients for 1) Caffeine, 2) interaction of Caffeine x A_1_R BP_ND_ in predicting the percentage of the identified cerebral responses on R (R core team, Vienna, Austria). The A_1_R BP_ND_ used was taken from the hemisphere corresponding to the detected cerebral responses.

Results are presented as mean ± standard deviation.

## Main results

### Salivary concentration

On CSR Day 5, the CAFF group had a significantly higher salivary caffeine concentration than the DECAF group (CAFF: 2.24 ± 1.16 mg/L, DECAF: 0.08 ± 0.07 mg/L, t_CAFF-DECAF_ = 9.9, p < .001). On REC Day, the salivary caffeine concentration in the CAFF group was lower (0.19 ± 0.31 mg/L, t_CSR-REC_ = 9.7, p < .001) and reached a level, which did not significantly differ from the DECAF group (0.01 ± 0.02 mg/L, t_CAFF-DECAF_ = 0.8, p = .871).

### Multimodal analyses on grey matter

The results of whole-brain analyses are illustrated in **Fig 2**, and the *t* values, *p* values, and Cohen’s *d* are detailed in **Table 1**. While no <u>main effect of CSR</u> on GM was found, the voxel-wise multimodal analysis controlled for the variances of CBF identified 7 large GM clusters which showed a <u>caffeine x CSR interaction</u>. The regions included: **Cluster 1**: *right* Rolandic operculum, **Cluster 2**: *right* thalamus, **Cluster 3**: *right* middle occipital gyrus, **Cluster 4**: *bilateral* medial superior frontal gyri and anterior cingulate cortex, **Cluster 5**: *left* sensory motor area and *bilateral* middle cingulate cortex, **Cluster 6**: *left* middle frontal gyrus, and **Cluster 7**: *left* precentral gyrus, inferior frontal operculum and insula.

**Figure 2.**
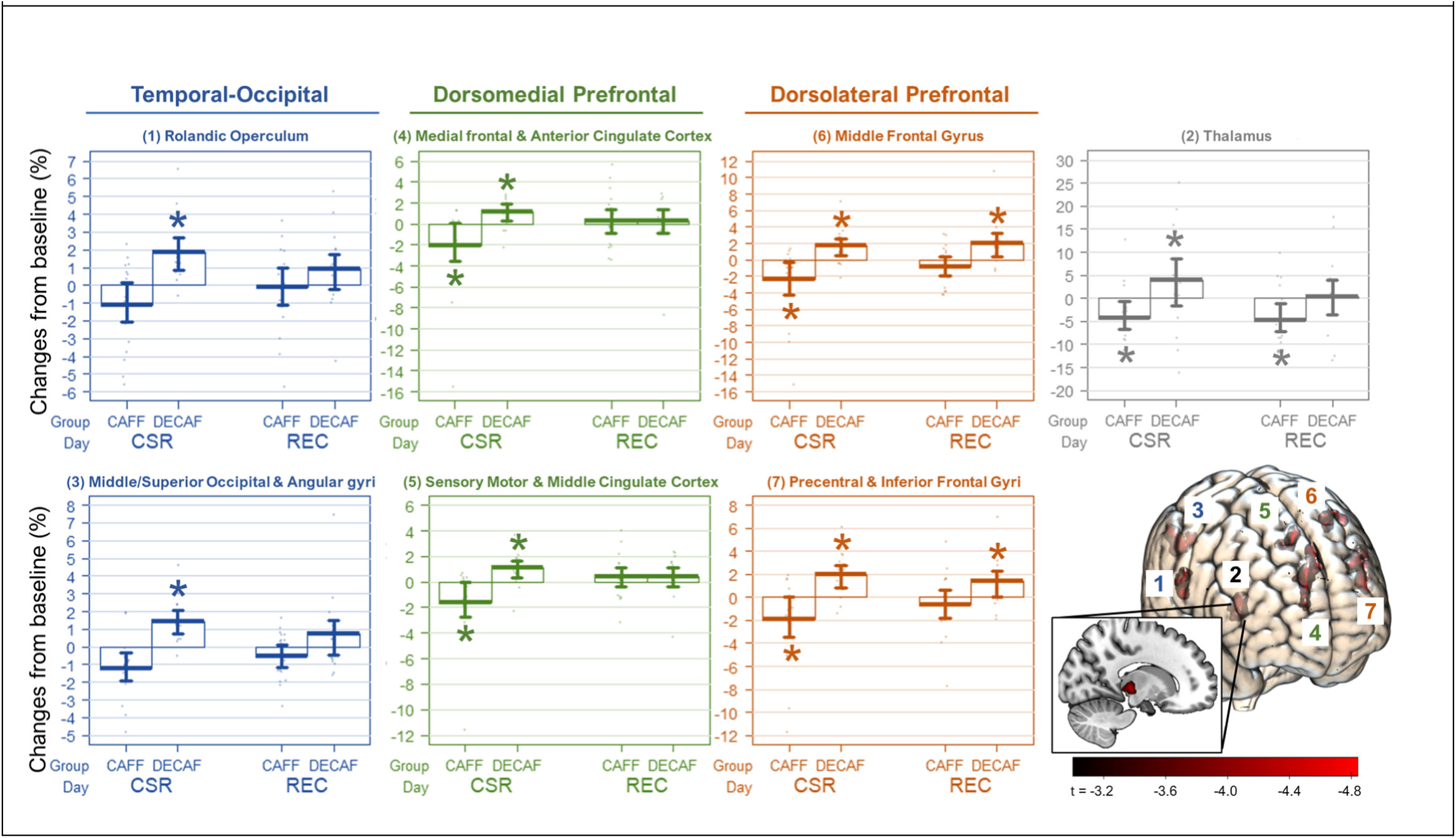
Regions showing significant differences between DECAF and CAFF groups in grey matter changes after CSR. The seven box plot panels display the clusters identified for a significant Caffeine x CSR interaction. The asterisks indicate a significant change compared to baseline as analyzed in post-hoc analyses. The color codes of the panels refer to a broader group of clusters based on functionality. The brain render visualizes the identified regions coded with the corresponding cluster number. The color bar indicates the t value acquired from the whole brain analysis. Finally, the render in multi-slices on the sagittal plane displays the exact location of the deeply seated Cluster 2.

**Table 1.** Post-hoc analyses on the changes in each GM cluster after CSR and REC compared to BL by groups (CAFF N=19; DECAF N=17). The d values refer to the effect size indicated by Cohen’d. The bold font indicates a statistical significance delineated by a p value < .05. The statistical parameters were controlled for age, sex, and total intracranial volumes in the linear mixed models. The definitions of regions are based on AAL2; voxel sizes in each region were estimated by MRIcron.

| Cluster | Regions (N of voxels) | Group | CSR – BL |  |  | REC – BL |  |  |
| --- | --- | --- | --- | --- | --- | --- | --- | --- |
|  |  |  | t | p | d [95% CI] | t | p | d [95% CI] |
| 1 | Rolandic Oper R (706) | **DECAF** | **3.9** | **<0.001** | 1.9 [0.7, 3.0] | 1.9 | 0.066 | 0.9 [-0.1, 1.9] |
|  | Postcentral R (135)<br>SMG R (65) | **CAFF** | -2.0 | 0.051 | -0.9 [-1.9, 0.0] | 0.0 | 0.971 | 0.0 [-0.9, 0.9] |
| 2 | Thalamus R (681) | **DECAF** | 1.9 | 0.062 | 0.9 [-0.1, 1.9] | 0.1 | 0.886 | 0.0 [-0.9, 1.0] |
|  | Hippocampus R (34)<br>Lingual R (34)<br>STG L (11) | **CAFF** | **-3.0** | **0.005** | **-1.4 [-2.4, -0.4]** | **-3.0** | **0.005** | **-1.4 [-2.4, -0.4]** |
| 3 | MOG R (827) | **DECAF** | **3.6** | **0.001** | **1.7 [0.6, 2.9]** | 2.0 | 0.056 | 1.0 [-0.0, 2.0] |
|  | SOG R (93)<br>AnG R (82) | **CAFF** | **-3.5** | **0.001** | **-1.6 [-2.6, -0.6]** | -0.8 | 0.423 | -0.4 [-1.3, 0.5] |
| 4 | mSFG L (812) | **DECAF** | **3.1** | **0.004** | **1.5 [0.4, 2.6]** | 1.1 | 0.259 | 0.5 [-0.4, 1.5] |
|  | mSFG R (105)<br>ACC L (191)<br>ACC R (584) | **CAFF** | **-2.7** | **0.009** | **-1.2 [-2.2, -0.3]** | 0.8 | 0.453 | 0.4 [-0.5, 1.3] |
| 5 | SMA L (375) | **DECAF** | **3.9** | **<0.001** | **1.9 [0.7, 3.0]** | 2.0 | 0.052 | 1.0 [-0.0, 2.0] |
|  | MCC L (336)<br>MCC R (186) | **CAFF** | **-3.1** | **0.003** | **-1.4 [-2.4, -0.4]** | 1.2 | 0.223 | 0.6 [-0.4, 1.5] |
| 6 | MFG L (506) | **DECAF** | **2.8** | **0.008** | **1.4 [0.3, 2.4]** | **3.2** | **0.003** | **1.6 [0.5, 2.6]** |
|  |  | **CAFF** | **-2.9** | **0.006** | **-1.3 [-2.3, -0.3]** | -0.8 | 0.421 | -0.4 [-1.3, 0.5] |
| 7 | Precentral L (784) | **DECAF** | **4.0** | **<0.001** | **1.9 [0.8, 3.1]** | **3.0** | **0.005** | **1.5 [0.4, 2.5]** |
|  | IFG Oper. L (1433) |  |  |  |  |  |  |  |
|  | IFG Tri. L (352) | **CAFF** | **-2.5** | **0.017** | **-1.1 [-2.1, -0.2]** | **-0.8** | **0.435** | **-0.4 [-1.3, 0.5]** |
|  | Rolandic Oper. L (256) |  |  |  |  |  |  |  |
|  | Postcentral L (123) |  |  |  |  |  |  |  |

**Abbreviations:** *R*, right hemisphere; *L*, left hemisphere. *ACC*, anterior cingulate cortex; *AnG*, angular gyrus; *IFG*, inferior frontal gyrus; *MCC*, middle cingulate cortex; *MFG*, middle frontal gyrus; *MOG*, middle occipital gyrus; *mSFG*, medial superior frontal gyrus; *Oper*, Operculum; *SMA*, supplementary motor area; *SMG*, supramarginal gyrus; *SOG*, superior occipital gyrus; *STG*, superior temporal gyrus; *Tri*, Triangulum.

Interestingly, the post-hoc analysis revealed similarities among the different clusters in their response to CSR and recovery sleep. On CSR Day 5 compared to BL, the DECAF group exhibited a GM increase in *all* clusters. The CAFF group, however, showed a GM reduction in all clusters except *right* temporal-occipital cortex (**Cluster 1 & 3**). On REC Day, the higher GM in the DECAF group had been remitted to the level of BL in *all* clusters except for *left* dorsolateral prefrontal cortex (DLPFC, **Cluster 6 & 7**), which in general crosses the bilateral anterior and middle cingulate cortices. Furthermore, the GM reduction in the CAFF group remitted in all clusters except for thalamus (**Cluster 2**). The response of **Cluster 2** GM inspired a supplementary correlation analysis between all the responsive regions (**Supplement**), on which the following A_1_R availability analysis (see paragraph *A1R BP_ND_ and GM plasticity*) was based.

### A_1_R BP_ND_ and grey matter plasticity

In order to reduce the bias from multiple comparisons, we synthesized GM responses into two groups – Cluster 2 (thalamus) and all others – based on the intercorrelation test between the Clusters responses (**Fig S2**). Overall, both cortical and subcortical A_1_R BP_ND_ could partly explain the group effect on the caffeine-associated GM reduction after CSR. However, subcortical A_1_R BP_ND_ showed a stronger and significant association with the caffeine-associated GM reduction after CSR, compared to cortical A_1_R BP_ND_. Furthermore, neither cortical nor subcortical A_1_R BP_ND_ showed significant association with the GM changes in Cluster 2 (thalamus). The associations of cerebral responses and A_1_R BP_ND_ are presented in **Fig 3**.

**Figure 3.**
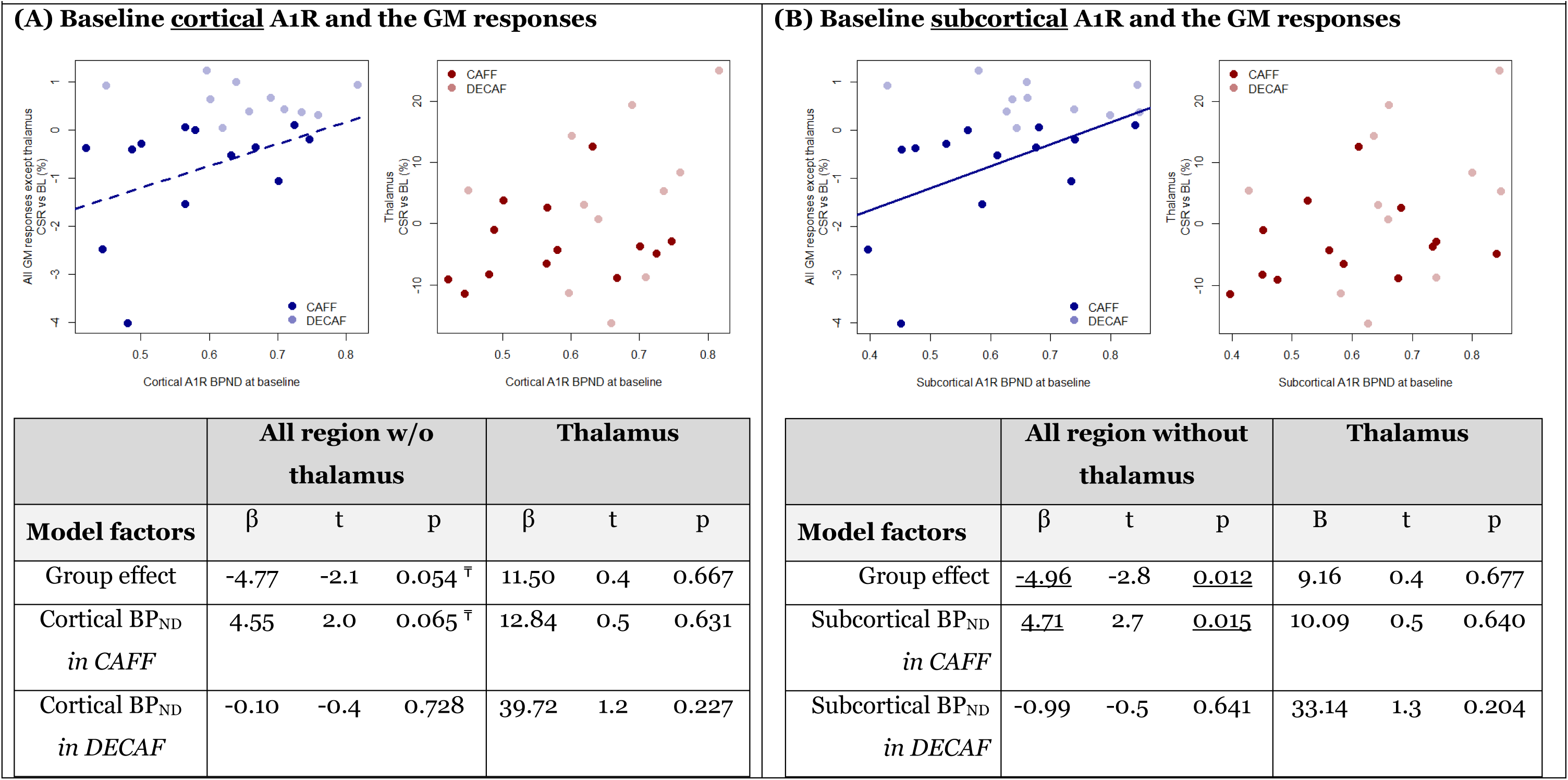
Associations between GM changes (% to baseline) and cortical or subcortical A_1_R BP_ND_. (A) and (B) display the associations of cortical and subcortical A_1_R BP_ND_ with GM responses, respectively. The statistical parameters are derived from analyses with linear mixed models. GM changes in all regions except thalamus are in blue while changes in thalamus are in red. Dark shades represent CAFF group while lighter shades represent DECAFF group. The solid line indicates a significant association between baseline A1R availability and the GM change, while a dash line indicates an association at trend. *<u>Table:</u>* Underscored p values were emphasized for the statistical significance, while ^₸^ mark refers to the effects that showed changes at trend.

### Exploratory analysis on cerebral blood flow

The whole-brain analysis on CBF did not indicate a significant difference between CSR Day 5 and BL Day or an interaction of caffeine with condition (i.e., CSR vs. BL)-BL. However, based on the solid evidence on the effects of caffeine and caffeine cessation on reducing and elevating CBF, respectively (9, 39–46), we examined the GM changes from CSR to REC, as well as a group difference in such changes. Interestingly, we found a significantly lower CBF on CSR compared to REC Day in the cerebellum, pons, brain stem, thalamus, hypothalamus, inferior temporal gyrus, and prefrontal cortex (all regions: p_FDR-corrected_ < 0.05; **Supplement Fig S1**); however, we did not find a significant group difference in the divergent responses between CSR and REC. To provide further information, we conducted a series of supplementary analyses on the effect of CSR or REC compared to BL Day with a lower statistical threshold specifically in the regions showing lower CBF on CSR Day than REC Day (i.e., the medial frontal cortex, subcortical regions, occipital cortex, cerebellum, and midbrain). We found a reduced CBF on the CSR Day and an elevated CBF on REC Day compared to the BL Day in the CAFF group. The statistics and figures are presented in the **Supplement**.

## Discussion

The current study examined the cortical plasticity in response to CSR with or without the concomitant interference of caffeine. Independent of the caffeine-induced variances in CBF, we found opposite GM responses between the caffeine and decaffeinated groups after CSR. Specifically, GM in the left DMPFC and DLPFC, as well as right temporal-occipital and thalamic cortices were increased after 5-day CSR without caffeine; in contrast, concomitant caffeine intakes during CSR led to a decrease in prefrontal and thalamic cortices. All the cortical changes, except for the DLPFC increase in the DECAF group and the thalamic decrease in the CAFF group, recovered to the baseline level after an 8-h sleep opportunity (as well as an approximately 35 h withdrawal from caffeine for the CAFF group). Most importantly, individuals with a lower availability of subcortical A_1_R at baseline showed a larger caffeine-associated GM decrease after CSR in all responsive regions except for thalamus. Taken together, our data reveal region-dependent GM plasticity after CSR, which diverges between the presence and absence of caffeine. Furthermore, individual traits in A1R availability may play a resilient role against the effects of caffeine on GM changes after CSR.

### Implications on use-dependent cortical and synaptic plasticity associated with adenosine systems

#### GM plasticity and synaptic strengths

Brain structural alterations induced by sleep loss have been frequently reported. Extensive sleep loss such as 36-h to 72-h sleep deprivation (7, 8) as well as one month of CSR (15) was found to lead to reduced GM or structural network. Moreover, 21-h extended wakefulness followed by 3h time in bed resulted in both reduced GM in thalamus, precuneus, and precentral gyrus as well as increased GM in insula (6). Our data showing increased GM in DLPFC, DMPFC, thalamus, and temporal-occipital regions after 5 days of CSR with 5-h time in bed daily seemingly conflicts to the GM reduction commonly observed. However, the inconsistency may lie at the total days of sleep restriction and the different amounts of sleep as daily recovery process. The synaptic homeostasis hypothesis (SHY) (47) elaborates how synaptic homeostasis may be maintained in sleep-wake cycles. It is suggested that the synaptic strengths are increased and saturated throughout wakefulness while downregulated as a restorative process during sleep (47). Extensive sleep loss, therefore, may lead to neuronal exhaustion and thereby the commonly observed GM reduction. However, partial sleep (i.e. 5-h time in bed) may allow partial brain restoration and enable the synaptic strengths to maintain its upregulated state below the extent of exhaustion for a limited amount of days with sleep restriction. Such an upregulated synaptic strengths might underlie the increased GM observed in this study. In addition, an earlier study examining the trajectory of GM changes throughout a 36-h sleep deprivation has found a non-linear course of responses (7). They observed an increased GM in striatum and cingulate cortices throughout the 36-h sleep deprivation. while other regional GM reductions only started to show at 32h onward such as in the thalamus, insula, and somatosensory association cortex (7). Their findings support the adaptability of brain structure to a limited extent of sleep loss through an upregulated state until exhaustion might occur. On the contrary, caffeine has been shown to reduce the synaptic long-term potentiation (LTP) in rodents (48) and LTP-like cortical activity in humans (49), which may counteract the sleep loss-induced cortical upregulation. In other words, the saturation and suppression of synaptic strengths might underlie the CSR-associated GM increase and caffeine-associated GM decrease, respectively.

#### Adenosine system in energy use-dependent synaptic and cortical plasticity

GM plasticity may occur in response to an increased or decreased need for energy resources, for which adenosine is a critical index as a byproduct of energy usage (20, 21). A study found a higher GM volume in the morning compared to the evening (50) and suggested that the time of day-dependent plasticity of GM may be an evolutionary adaption to the energy consumption of diurnality (50). In sleep homeostasis process, the elevated and reduced adenosinergic activity throughout wakefulness and sleep, respectively, is believed to be a biological index of the accumulation and dissipation of homeostatic sleep pressure (21). By blocking adenosine receptors, caffeine counteracts the increased sleepiness as one of the classic effects and switches on an “energy saving mode” by reducing cortical activity (51), cerebral metabolic rate of glucose (52), and cerebral metabolic rate of oxygen (CMRO2) (53, 54). Our data show that caffeine does not only counteract the CSR-associated GM increase but further downregulates the cortical intensity potentially through hindering the adenosine-mediated energy consumption. Our finding is in line with earlier evidence of a reduced or lower GM volume after 10-day 450 mg caffeine intake (9) or in habitual high-dose caffeine consumers (10) in various brain regions.

The association between striatal A_1_R availability and cortical reduction after CSR with caffeine implicates a role of adenosine system in the CSR– and caffeine-associated GM plasticity. Specifically, a higher A_1_R availability seems to exert a resilience function to caffeine-associated GM reduction. Such a A_1_R-dependent resilience has also been found to be against sleep deprivation-induced cognitive impairments (55). Interestingly, despite the cortical and subcortical A_1_R availability being highly correlated, we observe a much stronger association of caffeine-associated GM reduction with subcortical than cortical A_1_R availability. In striatum, A_1_Rs antagonistically interact with A_2A_R through a formation of G-protein coupled-receptor heterodimer as a “*concentration-dependent switch*” (56) that modulates presynaptic glutamate release based on the extracellular adenosine levels. In a regular extent of wakefulness, the tonic neuro-inhibition maintained by A_1_R is strengthened as the waking state lasts longer (**Fig 4**), leading to a reduced striatal presynaptic glutamate release and neural firing (57, 58) which is likely to underlie the lower GM in the evening compared to mornings (50). In relatively shorter e.g. 12h or 24h) sleep deprivation or sleep restriction (**Fig 4**), the rapidly elevated level of adenosine (22) triggers the phasic excitatory adenosine A_2A_ receptors (A_2A_R) which in turns inhibits A1R and enhances glutamate release (56, 59, 60) as well as neural plasticity (24, 61, 62), potentially leading to a GM increase as observed in our DECAF group after 5-day CSR. A prolonged neuroexcitation, e.g. by an extensive sleep loss such as a 36-h or 72-h deprivation or one-month sleep restriction might turn into a detrimental hyperexcitability (63), potentially underlying the reduced brain structure frequently reported (7, 8, 15). Caffeine, on the other hand, acutely enhances striatal glutamate release by its predominant antagonism at A_1_R (64); however, daily or chronic caffeine administration abolishes the effect of antagonism at A_1_R on increasing glutamate level (65), leading to the aforementioned suppressing effects on LTP and neuroplasticity (48, 49) and reduced GM (9, 10). Furthermore, through a gradually reduced affinity of A_2A_R to caffeine over daily or chronic intake (56, 60), a concomitant exposure to high extracellular adenosine level such during CSR allows the A_2A_R rebinding to endogenous adenosine even with the presence of caffeine, thereby regaining the inhibition to A1R through the A_1_R –A_2A_R heterodimers in striatum (56, 60). The association between striatal A_1_R availability and caffeine-associated GM reduction after CSR further implicate a role A_1_R may play in moderating the balance between the A_1_R –A_2A_R counteractions in synaptic signaling and GM plasticity after daily exposure to caffeine and sleep restriction.

**Figure 4.**
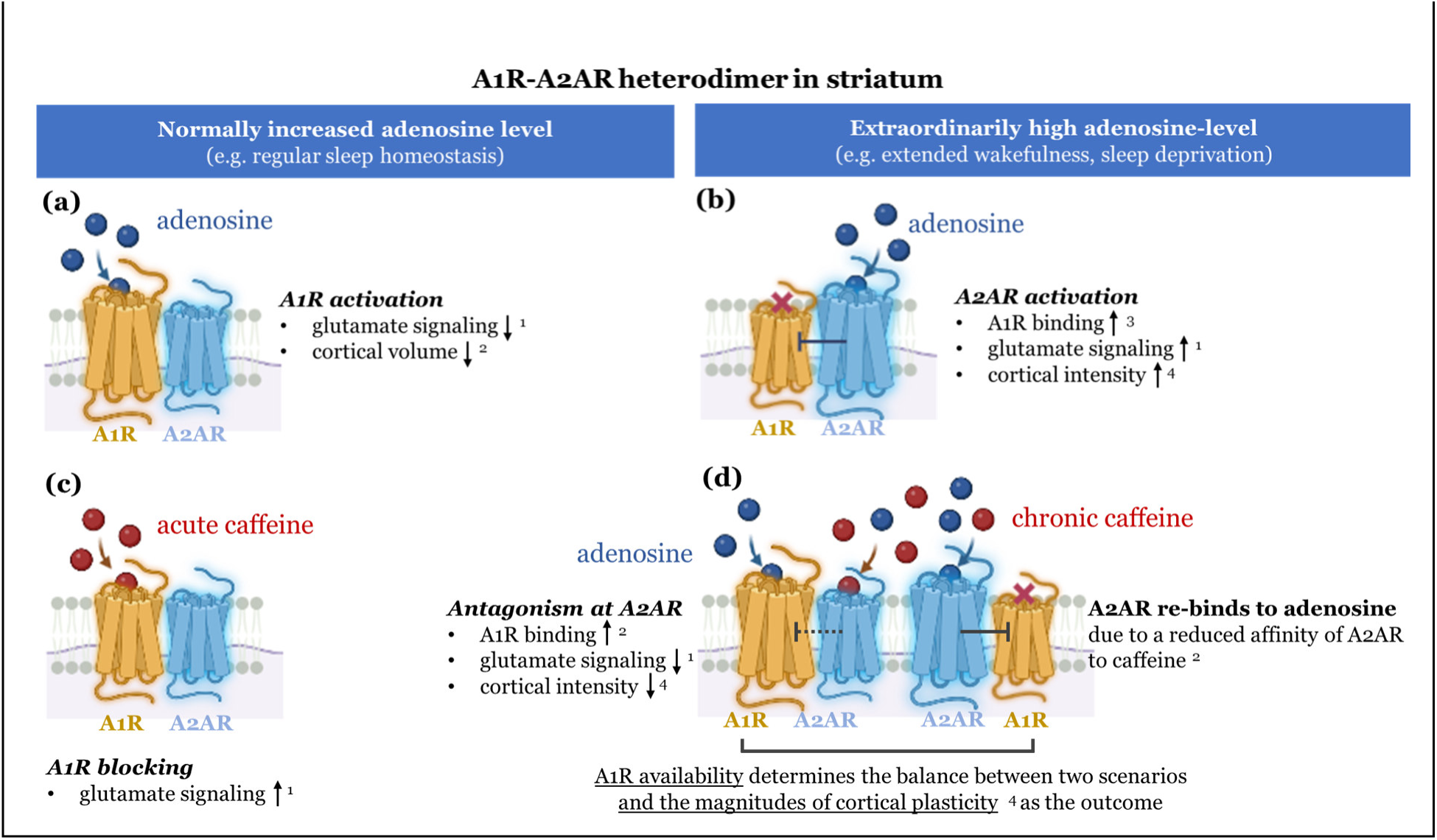
Schematic description of the A1R-A2AR interactions under different adenosinergic modulations and caffeine interference. In this schematic description, we summarize our results in the mechanistic context based on the literature (denotation: ^1^ Quarta et al. 2004 (68); ^2^ Trefler et al. 2014 (53); ^3^ Ciruela et al. 2006 (59)/ Ferre 2008 (63); ^4^ Current study observation). The left panel describes adenosine modulation under the circumstance of a regular sleep-wake cycle with (a) or without (c) caffeine administration. The right panel describes the adenosine modulation under the circumstance of extended wakefulness with (b) and (d) without caffeine administration.

### Beyond the use-dependent perspective: unique GM response in thalamus

Interestingly, although thalamic GM shows a consistent CSR-associated GM increase and caffeine-associated GM decrease, its recovery seems to be much slower than other responsive regions. An earlier study following up the trajectory of medial temporal GM recovery after the cessation of 10-day 450 mg caffeine intake suggested a fully recovery required longer than 36h (44). In this study, participants took a lower caffeine dose (i.e., 300 mg/day) in a shorter period of intake (i.e., 5 days), and the caffeine-associated GM reduction was nearly recovered in most of the responsive regions except for thalamus. Furthermore, the GM reduction in the thalamus is not directly associated with the individual A1R availability. An earlier study found regional differences in the change in adenosine signaling after sleep deprivation (66). While adenosine was rapidly accumulated and dissipated in basal forebrain and cortices after sleep deprivation and during recovery sleep, in thalamus, together with midbrain regions, showed a continuous decrease in adenosine concentration after sleep deprivation and even a few hours into recovery sleep (66). Such differences in adenosinergic activity between thalamus and other brain regions might underlie the unique responses of thalamus in our data. Since thalamus is one of the most critical regions in sleep-wake regulations (67, 68), and the structural or functional alterations of thalamus are often associated with insomnia (69–74) or hypersomnia (75), it may be impacted by the concomitant CSR and caffeine intake in a more critical manner compared to other cortical regions.

### Limitations and summary

A few limitations in this study require careful interpretation and further investigation in the future. First, although we preselected participants based on the *ADORA2A* polymorphism, the lack of A_2A_R availability data as derived from PET precludes a broader understanding of the receptor’s role in sleep loss induced GM changes. This limitation warrants more adenosine and A_2A_R-focusing studies on the effects of CSR and daily caffeine intake to be conducted. Second, it might be argued that the estimation of GM by MRI T1-weighted image can be potentially impacted by caffeine-induced changes in perfusion (30). However, our study included both structural imaging and CBF measurement, and we did not observe associations between the two, including the pattern of changes across regions. An earlier study further suggests that the covariance between T1-weighted measurement and CBF rather occurs on a global instead of regional level (9). Lastly, the caffeine and decaffeinated treatments in the current study were delivered as identically brewed coffee only different in the caffeine content, which indeed guaranteed the precision of caffeine administration. However, with other ingredients contained in brewed coffee, the observed outcomes could not be fully attributed to the effects of caffeine alone, but it could also be an effect from the interaction between caffeine and other biochemicals. Nevertheless, using brewed coffee is preferred given that it provides a better generalizability to a real-life coffee intake.

In summary, this study reveals reversible upregulated GM in frontal, temporal-occipital, and thalamic cortices in response to CSR. This plastic response, however, can be reversed and turned into a downregulated GM by concomitant caffeine intake. The individual baseline subcortical A_1_R availability may play a role in the caffeine-associated GM response. More studies are warranted to further investigate how the interaction between A_1_R and A_2A_R in mediating these GM changes.

