## Supplement for "Repeated Caffeine Intake Suppresses Cerebral Grey Matter Responses to Chronic Sleep Restriction in an A_1_ Adenosine Receptor-Dependent Manner"

**Figure S1. Supplementary analysis on the interaction effect of Caffeination x Day on CBF.** Panel (a): Regions that showed significant difference in CBF between CSR vs REC. The color bar refers to (1 – p value), i.e., the brighter the color is, the stronger the statistical significance is stated. Panel (b): the mean CBF response across the responsive region (CSR vs REC) on each day in each group. Supplementary analysis: In light of the contrast between CSR vs REC, we assumed an underpower issue and therefore attempted to provide more information on the trend of the CBF changes by lowering the statistical threshold ($P_{FDR\text{-}corrected} < 0.20$ but $> 0.05$). We expected to find a reduced CBF on the CSR Day and an increased CBF on the REC Day compared to BL. The voxel-wise whole-brain analysis with the reduced statistical threshold indicated an interaction effect between Caffeination x CSR in the regions corresponding to the contrast of CSR vs REC, i.e. the medial frontal cortex, subcortical regions, occipital cortex, cerebellum, and midbrain. The linear mixed model on the extracted CBF response indicated a significantly stronger reduction of CBF on the CSR ($t_{interaction} = -3.2$, $p_{interaction} = 0.002$) and elevation on REC Day ($t_{interaction} = 3.4$, $p_{interaction} = 0.001$) in the CAFF group compared to the DECAF group.

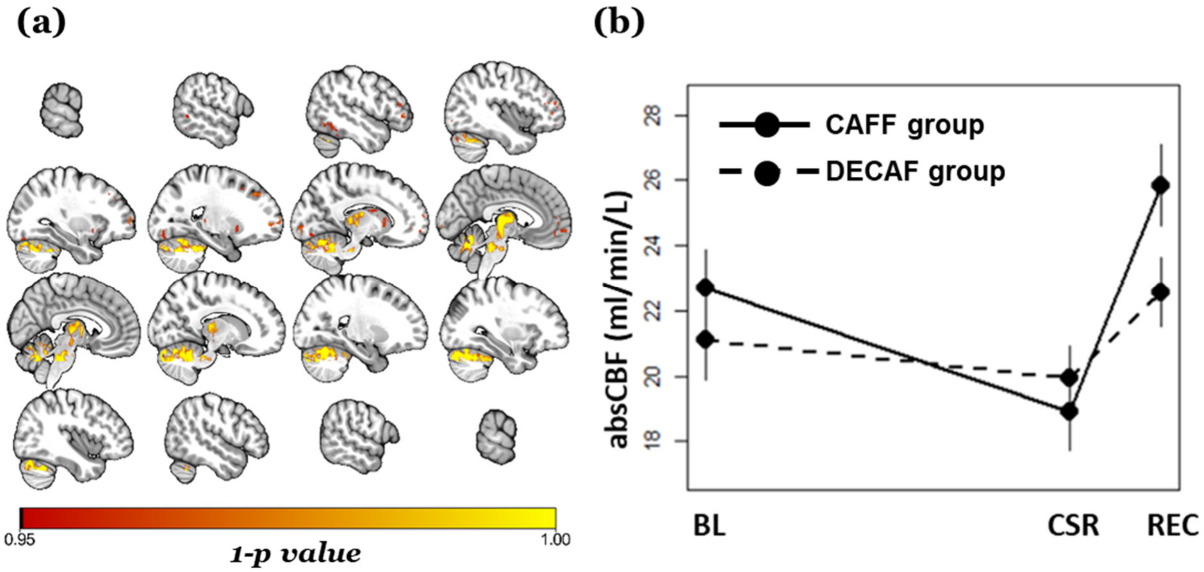

**Figure S2. Inter-correlation between Clusters.** The cross-correlation was analyzed and plotted using R package *corrplot (https://github.com/taiyun/corrplot)*. Panel (a) and (b) indicate the correlation between GM responses (i.e. changes from baseline) in CAFF and DECAF groups on CSR Day, respectively. The color bars below indicate the Pearson's correlation coefficient. The results of the cross-correlation suggest a uniquely weaker relationship between Cluster 2 (thalamus) and other clusters, and the discrepancy was particularly large in CAFF compared to DECAF group. This supplementary analysis was used to identify the similarities among GM responses and thereby group the regional data for further analysis.

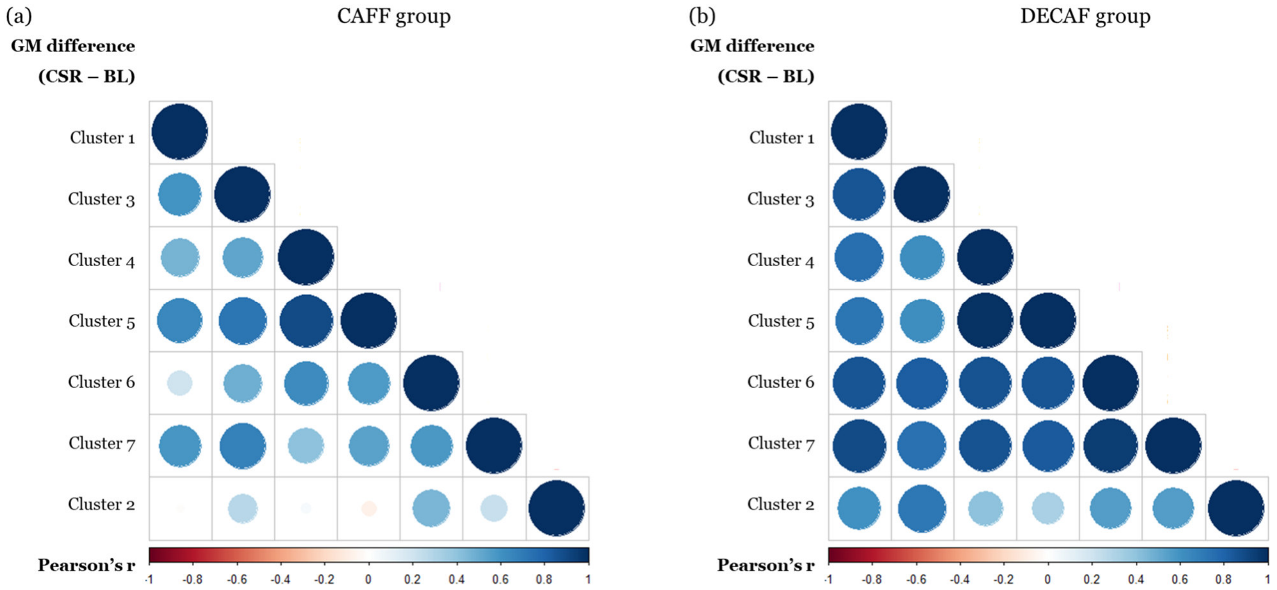
